# Streetlights affect moth orientation beyond flight-to-light behaviour

**DOI:** 10.1101/2022.10.06.511092

**Authors:** Jacqueline Degen, Mona Storms, Chengfa Benjamin Lee, Andreas Jechow, Anna Lisa Stöckl, Franz Hölker, Aryan Jakhar, Thomas Walter, Stefan Walter, Oliver Mitesser, Thomas Hovestadt, Tobias Degen

## Abstract

One of the most dramatic changes occurring on our planet in recent decades is the ever-increasing extensive use of artificial light at night, which drastically altered the environment nocturnal animals are adapted to ^1,2^. One nocturnal species group experiencing marked declines are moths, which are not only of great importance for species conservation, but also for their key role in food webs and in ecosystem services such as nocturnal plant pollination ^3,4^. Light pollution has been identified as a driver in the dramatic insect decline of the past years ^5–7^, yet little is known about its impact on natural insect orientation behaviour. Using harmonic radar tracking, we show that the orientation of several species of moths is significantly affected by streetlights, although only 4 % of individuals showed flight-to-light behaviour. We reveal a species-specific barrier effect of streetlights on lappet moths whenever the moon was not available as a natural celestial cue. Furthermore, streetlights increased the tortuosity of flight trajectories for both hawk moths and lappet moths. Our results provide the first spatially resolved experimental evidence for the fragmentation of landscapes by streetlights and demonstrate that light pollution affects movement patterns of moths beyond previously assumed extend, potentially affecting their reproductive success and hampering a vital ecosystem service.

## Introduction

The dramatic insect decline is one of the most concerning recent biological problems ^8,9^. Among insects, pollinators are of particular importance. Because of their significance for insect-pollinated plants, ecosystem functioning and food security, their decline will have severe implications for humans as well ^3,4^. While great focus has been dedicated to finding the causes and mitigating the decline of diurnal pollinators ^10,11^, nocturnal pollinator decline is less well understood. At night, moths belong to the most important pollinators ^12,13^ and there is also evidence for their decline in abundance and distribution ^14,15^. In addition to general drivers of insect decline ^16^, nocturnal pollinators are also threatened by light pollution ^5–7,17^. Light threatened by pollution ^18^ is caused by artificial light sources used privately and publicly, all of which differ from natural light sources in spectrum and intensity ^19^. Thus, artificial light at night (ALAN) changes and disturbs natural night environments with negative impacts from individual species to whole ecosystems, potentially affecting biodiversity ^20,21^. Furthermore, ALAN disrupts the natural visual cues nocturnal insects rely on for orientation ^22–24^. The most famous effect of artificial light sources is the strong phototactic response of moths, resulting in a flight towards light sources ^25,26^. Such “flight-to-light behaviour” has been the focus of the majority of investigations ^27^. Nevertheless, it is neither sufficiently understood why moths fly to the light, nor what exactly determines the attraction radius of a light source. Notably, as ALAN triggers maladaptive flight-to-light behaviour, it creates an “evolutionary trap” that reduces survival and reproduction ^28,29^. Because of methodological constraints, previous studies on the effects of streetlights were restricted to specific locations, using capture-recapture experiments ^30,31^ and observations within the light beam of a single lamp ^32^ or theoretical models ^33,34^. However, these results can only reveal the effects but not the causes for the impact of ALAN on moth behaviour. Understanding why streetlights affect movement behaviour and orientation performance requires measurements of the entire flight trajectories inside and outside of the illuminated area. We therefore used harmonic radar technology for the first time on several nocturnal pollinators, recording individual flight trajectories of moths at unprecedented spatial and temporal resolution within 1 km range.

## Results

### Hardly any moth terminated its flight at a streetlight

To investigate the influence of ALAN on the flight behaviour of moths, we recorded the flight trajectories of hawk moths (*Laothoe populi, Deilephila elpenor, Sphinx ligustri*) and lappet moths (*Euthrix potatoria*, Tab. S1) with harmonic radar (Fig. 1). Since this technique requires a certain handling procedure for the attachment of the necessary transponder, we confirmed in control experiments that flight behaviour was not significantly affected (see methods). Males were released one-by-one in the centre of six circularly arranged high pressure sodium streetlights (radius: 85 m, Fig. 1b). To compensate for daily fluctuations in weather and ambient light conditions, a similar number of individuals was tested each day with these lights either turned on or off. Out of the 50 animals that were released with the streetlights turned on, only two individuals (4%) terminated their flights directly at a streetlight (Fig. 1a, flight trajectories of the two individuals Fig. 1c). The positions of last waypoints of all other flights were widely scattered within the detection range of the radar (Fig. 1a) and there was no significant difference in the distance of the last recorded waypoint to the nearest streetlight between “light on” an “light off” conditions (Mann-Whitney *U*-test: *U*(50,45) = 1079.5, *P* = 0.735). To ensure that the light sources used in the experiments (Fig. S1 & S2) generally triggered the disrupted behaviour described in literature ^35^, we released seven moths of the species *Sphinx ligustri* in front of a streetlight at a distance of 10 m. All these males showed the typical behaviour of circling around the light at different heights and crashing to the ground from time to time until they stay motionless on the ground ^36^. This indicates that the streetlights we used influenced the behaviour in the expected, disruptive way within a close range (≤ 10 m) when the light source was above the moth at the time of release.

**Fig. 1.**
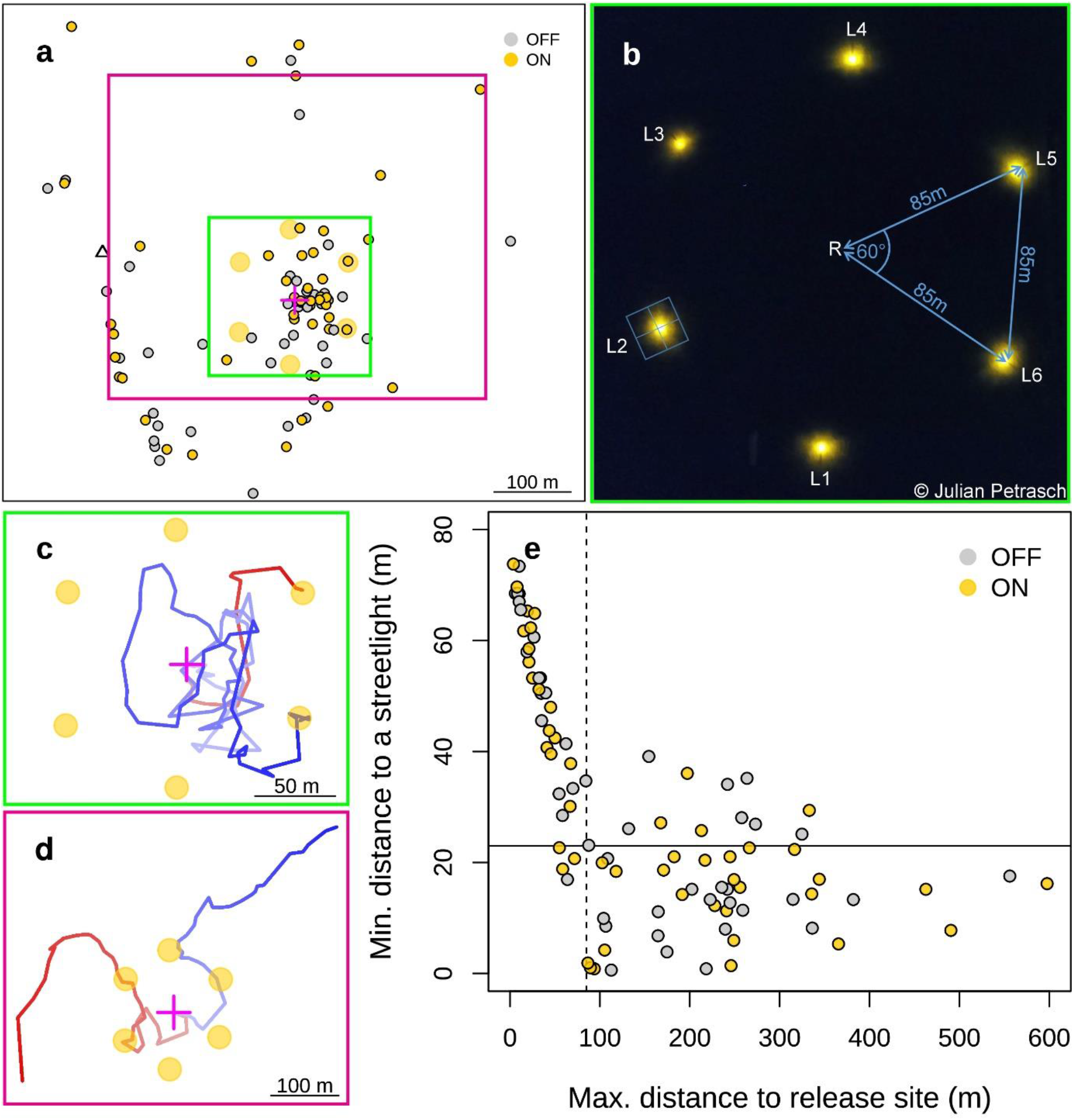
Final positions of tracked moths and flight proximity to a streetlight. **a**, Final recorded positions of all tracked moths (*n* = 95). **b**, Arrangement of the six high-pressure sodium streetlights used in the experiment imaged from a drone (picture taken by Julian Petrasch). The distance between the release site and each streetlight as well as the distance between them was 85 m. Please note that light cones of single streetlights did not overlap. Representative illuminance measurements of one streetlight (L2) are indicated by the blue square and corresponding values are illustrated in Fig. S1. **c**, Flight trajectories of the only two individuals that showed flight-to-light behaviour and ended their flight at a streetlight. **d**, Representative flight examples of individuals that passed an illuminated streetlight closer than 10 m and continued their flight (*n* = 6). **e**, Maximum distance to the release site and minimum distance to a streetlight at any time during a flight for all tracked moths (*n* = 95). The attraction radius of 23 m (indicated by the solid horizontal line) was calculated in a previous experiment using the same type of streetlights ^37^. Since each of the six streetlights was located 85 m away from the release site, this distance marks the minimum flight distance to arrive at a streetlight, as indicated by the dashed vertical line. **a-d**, All figures are aligned to the north.

Next, we analysed whether the males we released at the release site passed a streetlight within the attraction radius, i.e. the distance to a light that elicits flight-to-light behaviour, as males might have left the circle of the six streetlights without entering into any attraction radius. The attraction radius of high pressure sodium streetlights is estimated to be 23 m for moths in general ^37^. We therefore expected that all individuals that enter any streetlight’s attraction radius (Fig.1e, solid horizontal line) would show a positive phototactic response and thus terminate their flight near a light (Fig. 1e, dashed vertical line). However, apart from two moths that actually showed flight-to-light behaviour (Fig. 1c), 23 individuals entered the attraction radius of a streetlight but continued their flight and left the attraction radius again rather than showing flight-to-light behaviour (Fig. 1e, all individuals displayed with filled circles below the solid horizontal line and right of the dashed vertical line). The distance to streetlights passed during a flight in the “light on” and “light off” condition did not differ significantly (*t*-test: *t*_54_ = 0.434, *P* = 0.666) for moths that left the circle of streetlights (Fig. 1e, right of dashed line). We therefore conclude that most individuals were not attracted by the streetlights, even though they entered the attraction radius. Six moths (12 %) even passed an illuminated streetlight closer than 10 m without interrupting their flight (representative flight examples: Fig. 1d), a distance we have demonstrated to elicit flight-to-light behaviour when the animal was released from the ground (see above). Although the harmonic radar did not provide any information about the flight altitude, the flight direction could be communicated during the flight to the experimenter at the release site. This allowed to monitor the illuminated area of a streetlight more closely as soon as an individual approached it. Since we did not see any of the six individuals that passed a streetlight but continued their flight within the illuminated area, we hypothesise that they passed above the streetlight. We therefore suggest that flight altitude may be critical when assessing the attractiveness of a streetlight.

### Streetlights induced a species-specific barrier-effect

Although the light cones of the six circularly arranged streetlights did not overlap (Fig. 1b), this circle of streetlights might have created a barrier-effect, an “invisible wall” the moths were incapable to pass. Indeed, many individuals did not fly far enough to reach a streetlight (Fig. 1e, flights left of vertical dashed line) even though they initiated their flight properly and vanished from the field of view of the observer. Thus, these moths terminated their flight shortly after take-off. However, the streetlights did not create a barrier-effect for hawk moths, irrespective of the presence of the moon as a natural celestial cue (Fig. 2 a & b; Fisher tests for difference in fraction of animals leaving the circle with moon present and absent: *P* = 1, *P* = 0.57). In contrast, lappet moths were significantly prevented from leaving the circle of streetlights once these were turned on and the moon was not visible as natural celestial cue (Fig. 2 c & d, Fisher tests for difference in fraction of animals leaving the circle with moon present and absent: *P* = 0.58, *P* = 0.038). Thus, the illuminated circle of streetlights created a barrier effect for lappet moths when the moon was not visible. This is particularly interesting, because the wide stretches of unlit space between the streetlights (Fig. 1b) have not been sufficient enough for these moths to leave the circle of streetlights. Although we cannot determine which feature of moonlight enabled lappet moths to leave the illuminated circle of streetlights, the results confirm earlier findings showing that the moon can have a strong influence on the orientation of moths ^38^.

**Fig. 2.**
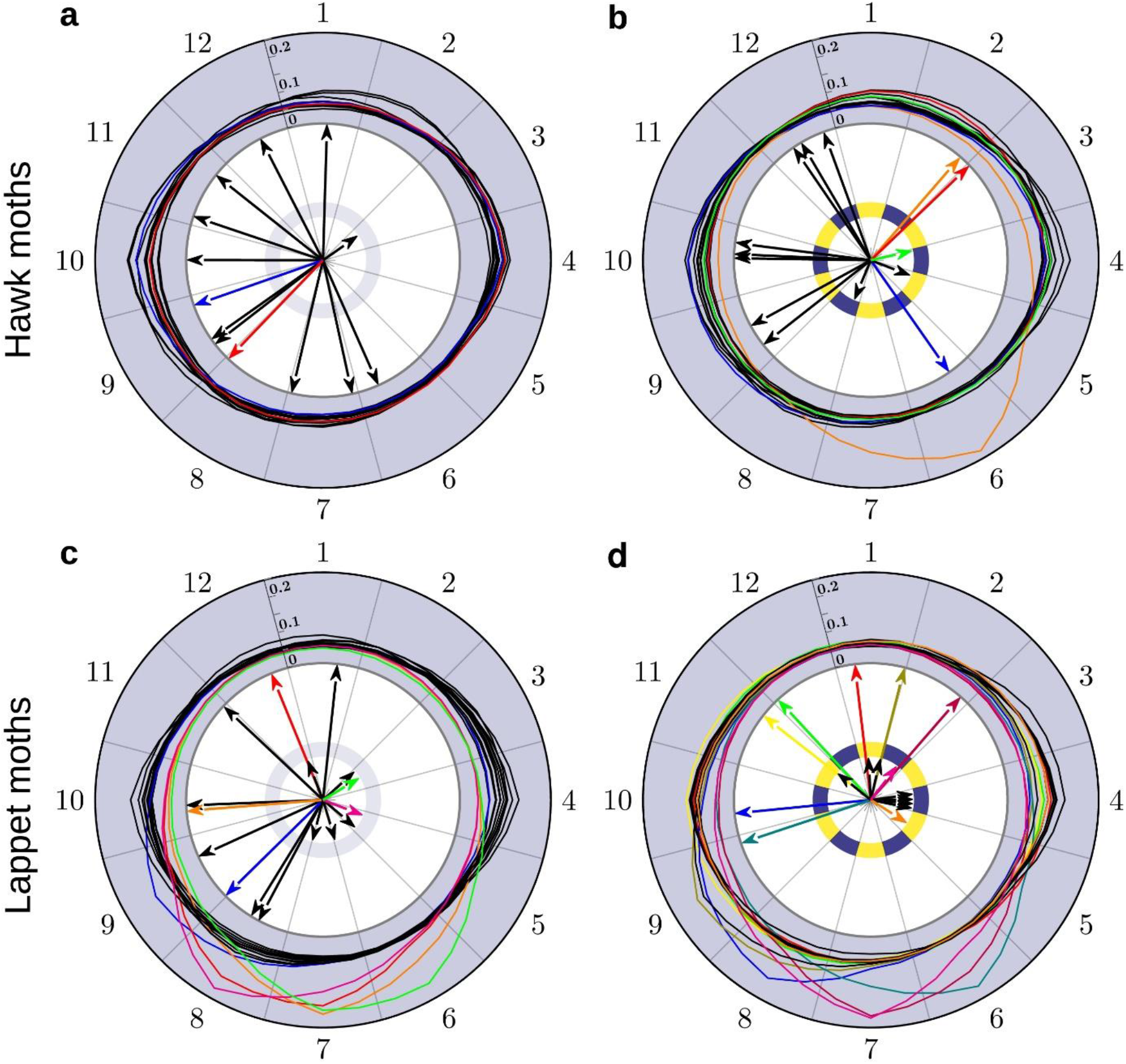
Linking flight directions and distances to the light environment. Flight directions were analysed separately for the conditions when lights were turned off (**a, c**) or on (**b, d**). For this analysis, the environment was divided into 12 sectors spanning 30° each, with odd-numbered sectors representing the position of a streetlight. The sectors are numbered clockwise in each plot, with the flight directions displayed as one arrow for each individual. Animals that did not leave the circle of streetlights are displayed with short arrows and those that left the circle by long ones. We divided all-sky images taken in parallel to the experiment (see methods) into the same sectors and calculated the mean luminance (“brightness”) for each single sector to link moth’s flight direction to the luminance of the surroundings. Luminance was normalized to compare light distribution patterns independent of varying light conditions of different nights (see methods) and the corresponding scale is displayed at the left boundary of sector one. Arrows when the moon was visible above the horizon are displayed in colour, matching the corresponding luminance distributions. Except when fully overcast (*n* = 4), the brightness was always highest in the sector where the moon was located, allowing to assess the flight direction in respect to the position of the moon. When the moon was below the horizon, flight directions as well as the corresponding brightness values are displayed in black.

Although we conducted the experiments in a relatively dark area, the surroundings featured a considerable amount of artificial light, ranging from streetlights of the close-by village Großseelheim to skyglow from distant cities (for details see methods). We quantified the light environment at the beginning of every flight via an all-sky image ^39^. Because the nocturnal light environment varied considerably between different nights, we normalized luminance for each image to identify the brightest sectors (Fig. 2). We found that the sectors with skyglow emerging from the towns Kirchhain and Stadtallendorf (Fig. 2, sector 4) and Marburg (Fig. 2, sector 10) were usually the brightest ones, with the moon overriding this pattern (e.g. Fig. 2b orange curve). Since flight directions were randomly distributed in all cases (Rayleigh-test for hawk moths with (*P* = 0.10) and without (*P* = 0.23), and for lappet moths with (*P* = 0.080) and without (*P* = 0.51) moon above the horizon), we conclude that moths did not fly into the direction of greatest sky brightness, respectively (weak) skyglow. This was also true when the moon was the brightest spot, indicating that the corresponding individuals (Fig. 2, flight directions (arrows) and brightness distribution (curves) are colour coded) did not fly directly in the direction of the moon.

### Streetlights increased the tortuosity of flights

The tortuosity of an animal’s path is a key parameter in orientation, including search behaviours, and is inversely related to the efficiency of the orientation mechanism involved for oriented flights while it reflects searching intensity for local search flights ^40^. Depending on the moths’ natural habitat, we expected different flight behaviours (directed or search flights) for different species. Since all hawk moth species in our study were collected outside of the experimental field, the test site can be assumed not to be a preferred habitat at this time of the year. We thus expected a directed and therefore straight flight out of the detection range of the radar to the surroundings. All lappet moths, on the other hand, already inhabited the experimental field and were expected to perform local search flights for resources (e.g. females). According to Benhamou ^40^, the tortuosity of flights needs to be calculated differently for oriented and search flights: while the tortuosity of oriented flights (hawk moths) needs to be calculated based on a straightness index, the tortuosity of local search flights (lappet moths) can be reliably estimated by a sinuosity index (see methods). To investigate the effect of streetlights on orientation and search behaviours, we therefore analysed whether turning on the streetlights elicited a change in the tortuosity of flights (Fig. 3).

**Fig. 3.**
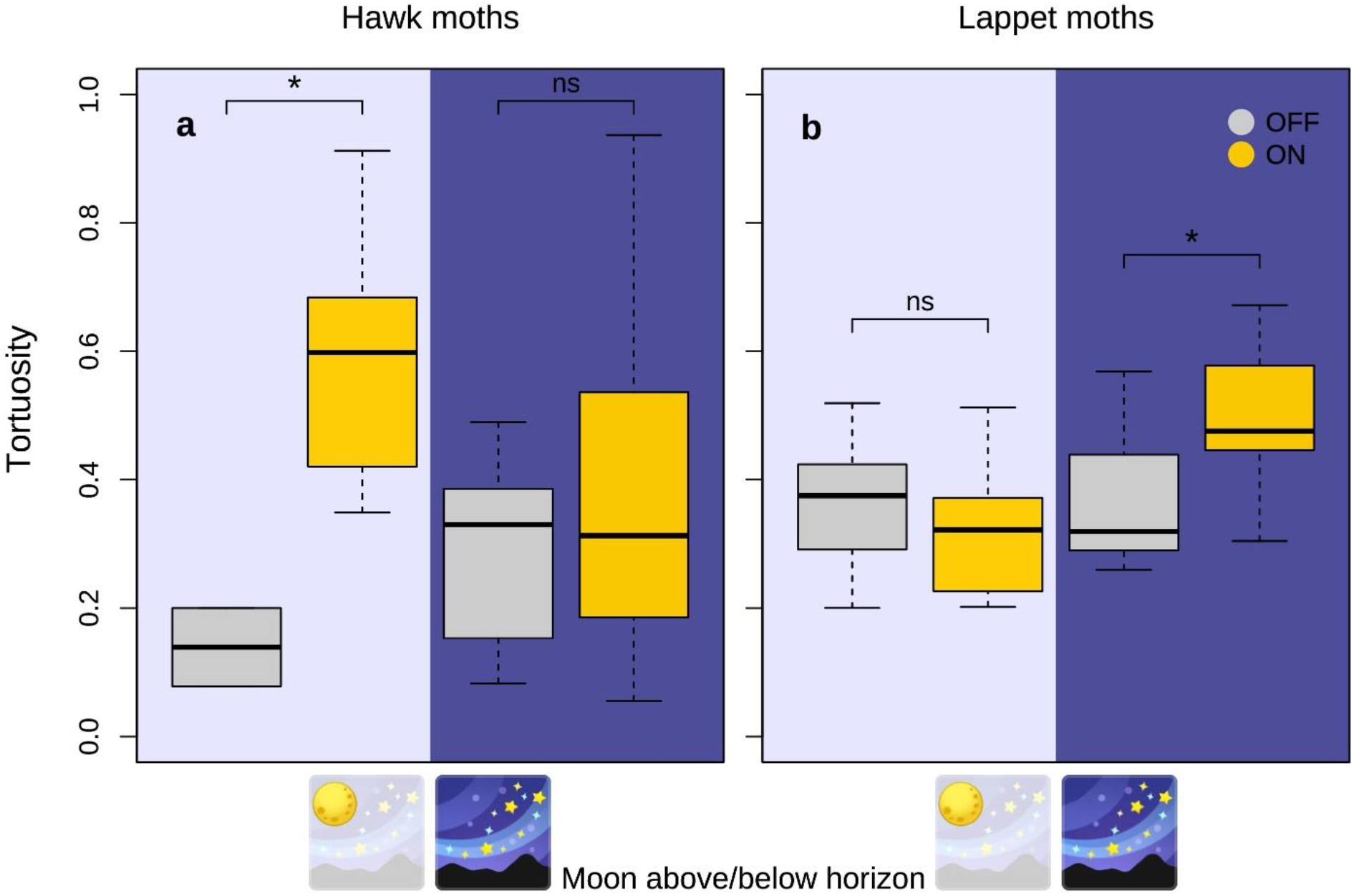
The effect of artificial light on the tortuosity of flights. **a, b**, Tortuosity of flights when streetlights were off or on in the presence or absence of the moon (sample sizes see Tab. S1). The tortuosity of oriented flights (hawk moths (a)) is inversely related to the efficiency of the orientation mechanism involved, while it reflects searching intensity for local search flights (lappet moths (b)). A value of 0 represents a perfectly straight flight and a value of 1 a very curvy flight. Values are displayed separately for hawk moths (**a**; *n* = 35) and lappet moths (**b**; *n* = 54), and nights when the moon was above the horizon (left) or below the horizon and therefore not visible (right). Box plots show the median (black line), the interquartile range (grey or orange box) and the minimum and maximum value within 1.5 times the interquartile range of the box (whiskers). Statistics: General Linear Model (GLM), significant differences (*P* < 0.05) are marked by *.

Hawk moths, which were not native to the experimental field, were expected to leave it fastest on a straight line. Indeed, when the streetlights were turned off, we observed rather straight flights, represented by a low tortuosity, especially when the moon was visible above the horizon (Fig. 3a, beta regression with post-hoc tests see Tab. S2). Switching on the streetlights significantly increased the tortuosity of flights when the moon was above the horizon, meaning that flights became less directed (Fig. 3a). Lappet moths, which were native to the experimental field, were expected to search for resources. Indeed, they generally had less directed flights compared to those of hawk moths when the streetlights were turned off, which likely reflects their search activity for local resources. When streetlights were turned on, the tortuosity of flights increased significantly when the moon was below the horizon (Fig. 3b, Beta regression with post-hoc tests see Tab. S3). Thus, our experiments revealed for both moth groups a significant change in flight behaviour when the streetlights were turned on.

### General discussion

The harmonic radar technique revealed a significant impact of streetlights on the flight behaviour of different species of moths even beyond the illuminated area. In addition to the barrier-effect on lappet moths, the significant increase in the tortuosity of flights caused by streetlights is of particular importance, because it relates to the orientation of individuals. Our results demonstrate for the first time that streetlights affect the orientation of moths although they do not show flight-to-light behaviour. This discovery adds a new dimension to the impact of light pollution on local movements of moths, which was previously not considered due to methodological constraints. Our finding that only very few moths showed flight-to-light behaviour although many entered the attraction radius of a streetlight raises the question why only such a low fraction got attracted. Generally, high pressure sodium streetlights are considered to be “insect-friendly” because of the spectral composition of their light emissions (Fig. S2, s.a. Eisenbeis ^36^), yet various studies have documented that nocturnal moths get attracted by and fly towards this type of lights ^41–43^. This is particularly true for hawk moths and lappet moths as demonstrated by light-trap catches ^37^. Our results suggest that the observation of moths trapped at streetlights only concern a small fraction of individuals that pass a streetlight in free-flight. Since we showed that orientation performance is negatively influenced by streetlights in general, light-trap catches might underestimate the impact of ALAN since only individuals showing flight-to-light behaviour are sampled. Although other explanations are possible, we emphasize the hypothesis that flight-to-light behaviour is triggered as a function of flight altitude, extending the attraction radius to a three-dimensional space. Thus, flight altitude might be of utter importance in this context and should be investigated in free-flying moths, using promising new methods that allow 3D-tracking once these have been fully developed for such demands ^44,45^.

The flight altitude of individuals may also explain why we found a barrier effect of streetlights for lappet moths but not for hawk moths (Fig. 2). Since the experimental lappet moths already inhabit the exact meadow where the experiments were performed while hawk moths do not, it seems reasonable to assume that lappet moths fly at lower altitudes to search for local resources while hawk moths may increase their flight altitude quickly after take-off to reach more favourable habitats. The barrier effect of streetlights on lappet moths is of particular importance, as it provides the first experimental evidence for the commonly postulated fragmentation of habitats by streetlights ^6,37,46^. Since the distance between the streetlights and thus the dark areas between the lights were unusually large compared to standard street lighting (Fig. 1b), it is likely that the barrier effect would be even stronger with the typical streetlight design. For example, in Europe pole distances of municipal streetlights for roads are between 25 and 45 m ^47^. Furthermore, we show a clear interaction between moonlight and ALAN, which should be taken into consideration for future studies on the impact of ALAN on nocturnal animals. Moon elevation and disk illumination should be reported in all studies, as effects of moonlight might mask or amplify the effects of ALAN.

Taken together, the harmonic radar technique revealed that streetlights affect moth orientation beyond flight-to-light behaviour, indicating a fundamentally novel dimension of impact at a local scale. This is of crucial importance for the probability of survival and mating success and supports the findings of Giavi et al. ^48^ that ALAN can affect ecosystem functioning in areas not directly illuminated. Since it has also been shown that ALAN is a thread to pollination ^49^ and potentially even alters diurnal plant-pollinator interactions ^50^, a reduced orientation performance of moths might represent a further parameter in fragile pollination networks. As the reduced orientation performance occurred independent of a disoriented behaviour caused by flight-to-light behaviour, we conclude that the negative effects of light pollution on moths have been underestimated to date.

## Methods

### Experimental Design

A harmonic radar (Raytheon Marine GmbH, Kiel, NSC 2525/7 XU) was used to track the flight paths of individual moths. This technique is well established for the investigation of navigation and orientation in honeybees ^51,52^, bumblebees ^53,54^ and diurnal pollinators ^55,56^ and could be easily conveyed to moths. The functional principle is described by Riley et al. ^57,58^.

The study site was located on an open flat pasture close to the small village Großseelheim, Germany. In the main experiment, all animals were released at the same location in the field (50°48’50.3”N, 8°52’32.7”E). Although the edge of Großseelheim was only about 430 m away from the release site and the towns Amöneburg, Kirchhain and Stadtallendorf (distance to release site: 3.7 km, 3.7 km and 10 km, respectively) as well as the cities Marburg and Giessen (Distance to release site: 7 km and 30 km) were not far away, the study area was relatively dark and not strongly impacted by skyglow ^38^. Six typical streetlights (GeoTechnik Kelvin-LED 1; c. 3.5 m high) equipped with high pressure sodium bulbs (70 W, 2000 K, 96 lm W^54^; NAV-E 70/E SON E27; Osram, Munich; Germany, s.a. Perkin et al. ^59^) were arranged uniformly in a circle around the release site with each light at a distance of 85 m to the release site and to its nearest neighbours (Fig. 1b). We used this type of streetlights to obtain representative results for the impact of common street lighting, since they are still one of most prevalent types ^60^. The lights were either switched off to record the flight trajectories under conditions without near-by artificial lights, or switched on to test the influence of streetlights on flight behaviour. It is important to note that the light cones of the lights did not overlap (Fig. 1b).

The experiments were performed from 10 June 2018 until 29 July 2018. During this time, we recorded 95 flights of 94 individuals of various species, nearly all of them either belonging to the family of lappet moths or hawk moths (Tab. S1). All hawk moths were collected with a large light trap that was built up every night at changing locations in the surroundings of the experimental area, far enough away to exclude visibility from the release site. Lappet moths were captured at the experimental field before the start of experiments. To this end, field paths were slowly followed with a car. Once a lappet moth got into the spotlight of the car, it merely made uncoordinated movements on the ground and could be captured easily.

After a moth was captured, it was kept in the dark and transported to the release site. Between capture and release of a moth there was a minimum acclimation time of 40 min (usually more than 60 min), and we assume that animals were dark-adapted at the time of take-off. When the animals were kept for longer times, they were fed with sugar solution (2M) to ensure that they had enough energy to perform a flight (except for *Euthrix potatoria* that do not assimilate food as adults). The animals were prepared with the transponder, the necessary antenna for radar tracking, shortly before their release. The procedure to attach the transponder to the thorax of a moth takes about 30 s and requires some light. To ensure that the moths’ vision did not get affected, we used only red light, which is not perceivable by most moth species including Sphingidae ^61^. Additionally, we tested a possible impact of the handling procedure, including the use of red light, during the control experiments (see below). We were able to follow the animals’ flights for up to 1 km with the position updated every 3 s.

### Light environment

Moon phase and position were retrieved from https://www.timeanddate.de. The nocturnal light conditions were monitored with a calibrated all-sky camera (Canon EOS 6D, Sigma EX DG 8mm fisheye lens 180°) ^38,39,62^. By obtaining an image at the start of each flight, we were able to measure spatially resolved sky brightness for each flight. For the analysis, luminance (L_v_ unit mcd/m^2^) was calculated for each pixel with the software “Sky Quality Camera” (version 1.8.1, Euromix, Ljubljana, Slovenia).

Illuminance and spectra of each streetlight were measured with a spectroradiometer in irradiance mode with a cosine corrected detector head (JETI Specbos 1211UV, Jena Technische Instrumente, Jena, Germany) at a height of 1.5 m because the vegetation did not allow a measurement exactly at ground level. Illuminance measurements were performed in a grid using a 2 m spacing along the main axis of the streetlight up to a distance of 10 m. Outer grid points were obtained in a 5 m spacing. An example grid is shown in Fig. S1 and an example spectrum in Fig. S2. Apart from the main axes, we measured at intervals of 5 m. The illuminance measurements and the drone image obtained at the beginning of the experiment revealed that lamp 3 (L3) had to be replaced to ensure equal brightness for all six lamps.

### Control experiments

To assess possible effects of the preparations needed for flight tracking via harmonic radar on natural flight behaviour ^63,64^, we performed four different control experiments with other males of the species *Sphinx ligustri* than those tested during the experiment with the harmonic radar. To this end, males were released from the same release site as the ones of the radar experiment, but the six streetlights were not turned on at any time. To create goals in the field, females (also *Sphinx ligustri*) operating as pheromone traps were positioned north and south of the release site in a distance of 105 m. We were therefore able to record the arrival frequency as well as the time males needed to reach the females using a stop watch. The same males were (1) prepared with a transponder and fed with sugar solution (2M) more than three hours before they were released. Afterwards, they were stored on a little wooden plate below a tin until the start of experiments, allowing a release without the need of the handling procedure to attach the transponder or the use of any light. On another day, these males were (2) prepared with a transponder directly before the flight, (3) experienced the same handling procedure as the animals in (2) but without attaching a transponder and (4) were released without a transponder and experienced no handling procedure at all by just storing them below tins as in experiment (1). Thus, the same set of males was tested in all four experiments, but not necessarily every individual went through all four experiments. Neither the arrival frequency (Fisher’s exact test: *P* = 0.887), nor the time successful males needed to reach the females located 105 m away differed significantly between the four groups (GLMM: *F*_3,31_ = 0.81, *P* = 0.505). In accordance with our former results acquired for honeybees ^65^, we can therefore be confident that the flight behaviour was not significantly affected by the tracking technique in our experiments.

### Data analysis

For the detailed analysis of flight behaviour (Fig. 2 & 3), only hawk moths and lappet moths were analysed due to sample size (Tab. S1). Flights with a total flight distance below 85 m that could not have reached a streetlight or with less than five recorded waypoints were not included in this dataset. To investigate the local impact of the streetlights we added to the experimental field, we analysed flight trajectories up to a distance of 270 m from the release site as this was the maximal possible tracking range in the direction of the village Großseelheim for safety reasons. For the evaluation of the main flight direction displayed with arrows in Fig. 2, we determined the mean cardinal direction from the release site for every flight ^66^. Hawk moths and lappet moths were not analysed together because they are native to different habitats and therefore perform different kinds of flights. Since hawk moths were not native to the experimental field, they should perform oriented and therefore rather straight flights to reach a more favourable habitat as fast as possible while lappet moths that are native to the experimental field should perform search flights to localize resources (e.g. females). This is especially relevant for the calculation of the tortuosity (Fig. 3), because a search path for local resources (lappet moths) differs from oriented flights to other landscape patches (hawk moths). According to Benhamou ^40^, tortuosity was therefore analysed by calculating a sinuosity index for lappet moths and the straightness for hawk moths using a special R package (R package trajr ^67^). For the calculation of the straightness, the distance of the beeline corresponded to 270 m for all individuals that left the radius of analysis (see above). For individuals that did not leave this circle, the beeline was calculated by subtracting the distance between the first waypoint of the trajectory to the release site from the distance between the last waypoint to the release site.

The software “Sky Quality Camera” (latest version 1.8.1, Euromix, Ljubljana, Slovenia) was used to calculate luminance values of 12 sectors spanning 30° each for the all-sky images. Since light conditions varied considerably between different nights, luminance values were normalized to compare light distribution patterns of different nights (Fig. 2). To normalize the values of the sectors, the mean luminance of the entire image was used:

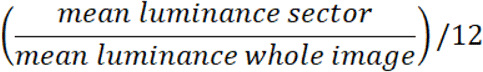

Consequently, normalized values reflect the contribution of each sector to the mean overall luminance. Thus, the sum of all 12 segments equals the total contribution (100 %) to the mean overall luminance of an all-sky image.

### Statistics

We utilized U-tests (*t*-tests) to analyse differences between lights on and off conditions for the distance of last recorded waypoint to the closest light source as well as closest distance during flight to any light. To test for differences between the four control experiments we used Fisher’s exact test to analyse arrival frequency and a Generalized Linear Mixed Model (GLMM) with the experiment as fixed effect and the individual as random effect to analyse flight duration. All statistical tests specified so far have been conducted with SPSS (IBM SPSS Statistics Version 26), all those mentioned in the following with R ^68^. Rayleigh tests for deviation from uniform circular angle distributions allowed identification of directional preferences (R package CircStats 0.2-6 ^69^) and Fisher’s exact tests identified differences in final positions inside vs. outside the lamp circle. Beta regression with Tukey post-hoc tests revealed differences in tortuosity of flights (R package glmmTMB 1.0.2.1 ^70^).

## Ethical Note

Our study involved individuals of several moth species (Tab. S1) that were trapped in the wild. We obtained permission for capture and release from the Regional Council of Giessen, Germany. All moths were carefully handled during experiments and maintained under appropriate conditions.

## Supplementary Information

Supplementary Information includes three tables and two figures.

## Acknowledgements

We thank Professor Menzel for the possibility to rent the harmonic radar for our experiments and the farmers in Großseelheim who gave us access to their grassland. Furthermore, we would like to thank Julian Petrasch for capturing our setup from above with a drone. Funding was provided by DFG DE 2869/1-1.

## Author contributions

J.D. designed the study, and M.S., A.Ja., S.W., T.W. and J.D. performed the experiments with substantial contributions of A.L.S. Radar data were extracted by M.S. and J.D., and M.S. and J.D. acquired and evaluated the data to quantify the light environment with substantial contributions of A.Je. and F.H. Data were analysed by T.D., O.M., C.B.L., T.H. and J.D. The original drafting of the article was done by J.D. and all authors contributed to the editing of subsequent drafts.

