## Supplementary Information for "Streetlights affect moth orientation beyond flight-to-light behaviour"

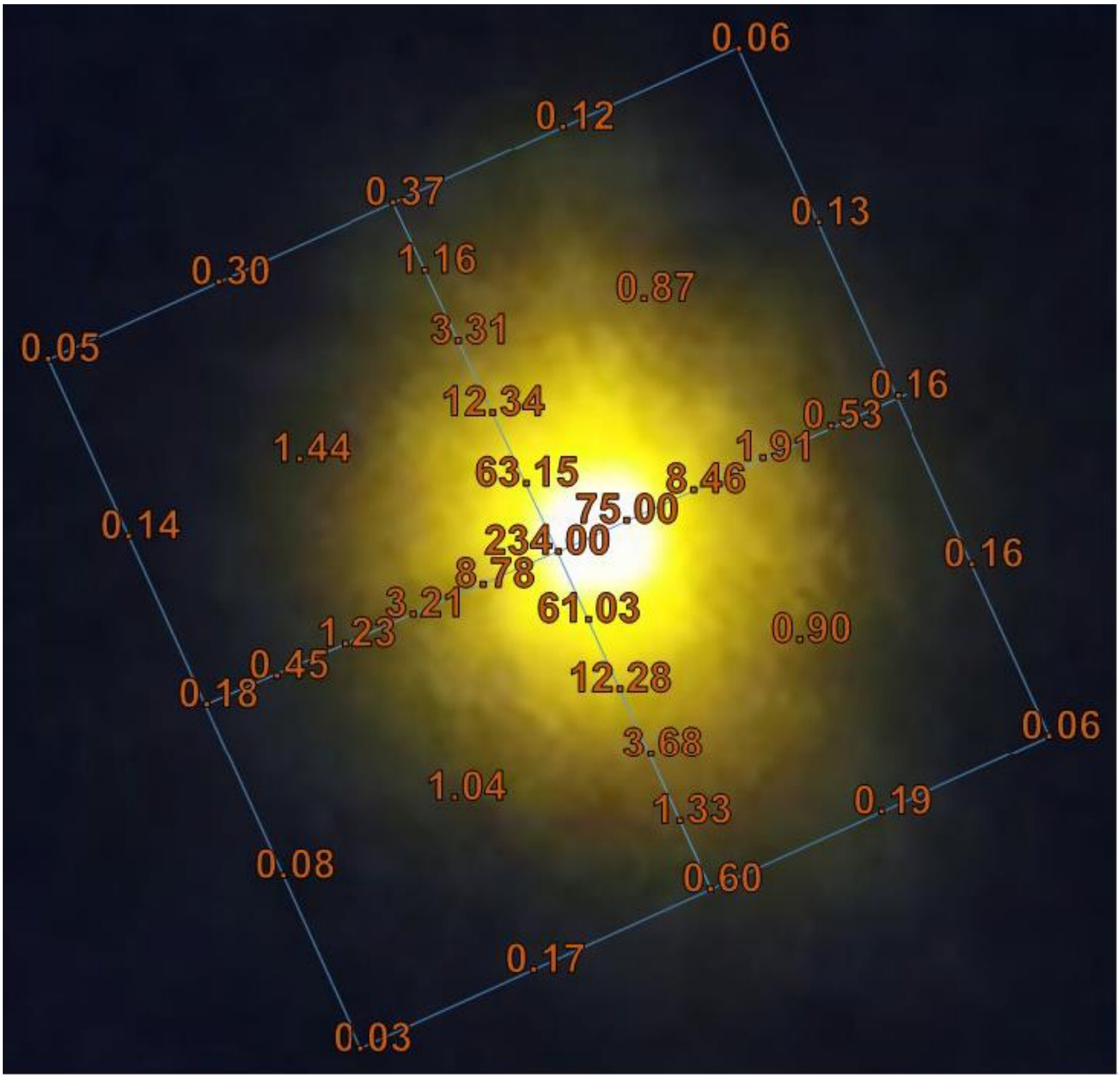


**Fig. S1 Representative illuminance measurement of a high-pressure sodium streetlight.** Illuminance (in lx) measured at a height of 1.5 m overlaid with a zoom in of the drone image (picture taken by Julian Petrasch) shown in Fig. 1b. The edge length of the blue square is 10 m.

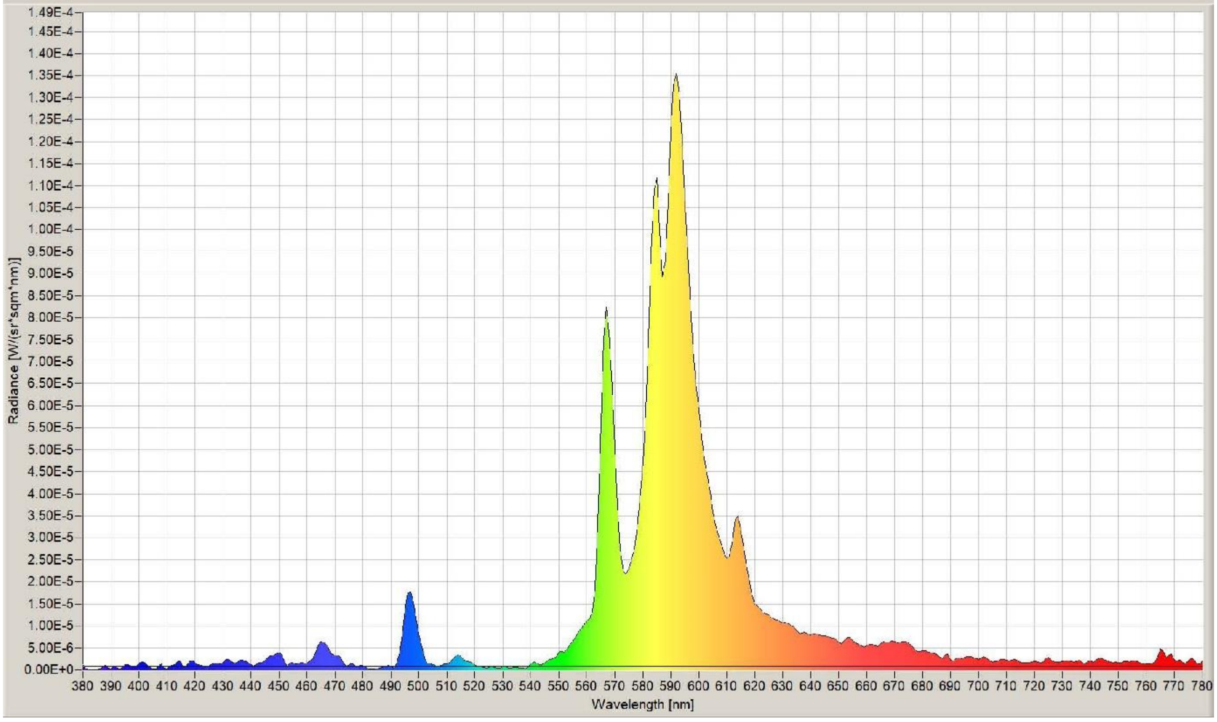


**Fig. S2 Spectrum of the high pressure sodium streetlights.** For the measurements, we used a spectroradiometer in irradiance mode with a cosine corrected detector head (JETI Specbos 1211UV, Jena Technische Instrumente, Jena, Germany) at a height of 1.5m.

**Tab. S1 Number of recorded flights.** Numbers are given for the different species with the six high-pressure sodium streetlights either turned off or on and the moon above or below the horizon.

| Family | Species | Recorded Flights | | | |
|---|---|---|---|---|---|
| | | Lights off | | Lights on | |
| | | Moon | No moon | Moon | No moon |
| Hawk moths | *Laothoe populi* | 1 | 9 | 4 | 6 |
| | *Deilephila elpenor* | 0 | 5 | 2 | 2 |
| | *Sphinx ligustri* | 1 | 3 | 0 | 2 |
| Lappet moths | *Euthrix potatoria* | 5 | 20 | 13 | 15 |
| Others | | 1 | 0 | 2 | 4 |

**Tab. S2: Beta regression of tortuosity in hawk moths.** Interaction: Lights on and moon below horizon.

| | Estimate | Std. Error | z value | Pr(>\|z\|) |
|---|---|---|---|---|
| (Intercept) | 1.410 | 0.662 | 2.128 | 0.033 * |
| Lights: On | -1.817 | 0.766 | -2.371 | 0.018 * |
| Moon: Below horizon | -0.597 | 0.701 | -0.851 | 0.395 |
| Interaction (Light, Moon) | 1.482 | 0.842 | 1.760 | 0.078 |

step = over: Light On vs. Off: $P = 0.025$

step = under: Light On vs. Off: $P = 0.35$

**Tab. S3: Beta regression of tortuosity in lappet moths.** Interaction: Lights on and moon below horizon.

| | Estimate | Std. Error | z value | Pr(>\|z\|) |
|---|---|---|---|---|
| (Intercept) | -0.575 | 0.194 | -2.966 | 0.003 ** |
| Lights: On | -0.171 | 0.244 | -0.701 | 0.483 |
| Moon: Below horizon | 0.049 | 0.233 | 0.210 | 0.834 |
| Interaction (Light, Moon) | 0.696 | 0.318 | 2.189 | 0.029 * |

step = over: On vs. Off: $P = 0.49$

step = under: On vs. Off: $P = 0.016$
